# High Aspect Ratio Nanomaterials Enable Delivery of Functional Genetic Material Without DNA Integration in Mature Plants

**DOI:** 10.1101/179549

**Authors:** Gozde S. Demirer, Huan Zhang, Juliana L. Matos, Natalie Goh, Francis Cunningham, Younghun Sung, Roger Chang, Abhishek J. Aditham, Linda Chio, Myeong-Je Cho, Brian Staskawicz, Markita P. Landry

## Abstract

Genetic engineering of plants is at the core of sustainability efforts, natural product synthesis, and agricultural crop engineering. The plant cell wall is a barrier that limits the ease and throughput with which exogenous biomolecules can be delivered to plants. Current delivery methods either suffer from host range limitations, low transformation efficiencies, tissue damage, or unavoidable DNA integration into the host genome. Here, we demonstrate efficient diffusion-based biomolecule delivery into tissues and organs of intact plants of several species with a suite of pristine and chemically-functionalized high aspect ratio nanomaterials. Efficient DNA delivery and strong protein expression without transgene integration is accomplished in *Nicotiana benthamiana* (*Nb*), *Eruca sativa* (arugula), *Triticum aestivum* (wheat) and *Gossypium hirsutum* (cotton) leaves and arugula protoplasts. We also demonstrate a second nanoparticle-based strategy in which small interfering RNA (siRNA) is delivered to *Nb* leaves and silence a gene with 95% efficiency. We find that nanomaterials not only facilitate biomolecule transport into plant cells but also protect polynucleotides from nuclease degradation. Our work provides a tool for species-independent and passive delivery of genetic material, without transgene integration, into plant cells for diverse biotechnology applications.

---

Plant biotechnology is critical to address the world’s leading challenges in meeting our growing food and energy demands, and as a tool for scalable pharmaceutical manufacturing. In agriculture, genetic enhancement of plants can be employed to create crops that have higher yields and are resistant to herbicides^1^, insects^2^, diseases^3^, and abiotic stress.^4^ In pharmaceuticals and therapeutics, genetically engineered plants can be used to synthesize valuable small-molecule drugs and recombinant proteins^5^. Furthermore, bioengineered plants may provide cleaner and more efficient biofuels^6,7^.

Despite several decades of advancements in biotechnology, most plant species remain difficult to genetically transform^8^. A significant bottleneck facing efficient plant genetic transformation is biomolecule delivery into plant cells through the rigid and multi-layered cell wall. Currently, few delivery tools exist that can transfer biomolecules into plant cells, each with considerable limitations. *Agrobacterium-mediated* delivery^9^ is the most commonly used tool for gene delivery into plants with limitations of efficient delivery to a narrow range of plant species and tissue types, inability to perform DNA- and transgene-free editing^10^, and unsuitability for high-throughput applications. The one other commonly used tool for plant transformation is biolistic particle delivery (also called gene gun)^11^, which can deliver biomolecules into a wider range of plant species but faces limitations of only bombarded-site expression, plant tissue damage when high bombardment pressures are used^8^, possible limitation of the specimen size and positioning in the biolistic chamber, and the requirement of using substantial amount of DNA to achieve desired delivery efficiency. To-date, there has yet to be a plant transformation method that enables high-efficiency gene delivery, without transgene integration, in a plant species-independent manner. Herein, we address the long-standing challenge of biomolecule delivery to mature model and non-model plants with nanomaterials, filling a key void in the plant transformation toolkit.

While nanomaterials have been studied for gene delivery into animal cells^12,13^, their potential for plant systems remains under-studied^14^. Under certain surface chemistries, high aspect ratio nanomaterials such as carbon nanotubes (CNTs) have recently been shown to traverse extracted chloroplast^15^ and plant membranes^16^ with several figures of merit: high aspect ratio, exceptional tensile strength, high surface area-to-volume ratio, and biocompatibility. When bound to CNTs, biomolecules are protected from cellular metabolism and degradation^17^, exhibiting superior biostability compared to free biomolecules. Moreover, single-walled carbon nanotubes (SWCNTs) have strong intrinsic near-infrared (nIR) fluorescence^18,19^ within the tissue-transparency window and thus benefit from reduced photon scattering, allowing for tracking of cargo-nanoparticle complexes deep in plant tissues. However, previous incorporation of CNTs in plant systems is limited to exploratory studies of CNT biocompatibility^15,20,21^ and sensing of small molecules in plant tissues^16,22^ by introducing CNTs complexed to synthetic fluorescent dyes or polymers.

Herein, we develop a CNT-based platform that can deliver functional biomolecules into both model and crop plants with high efficiency, no toxicity, and without transgene integration: a combination of features that is not attainable with any plant transformation approach. We used covalently-functionalized or pristine CNTs to deliver DNA into mature *Nb*, arugula, wheat, and cotton leaves, and obtained strong protein expression. We also show CNT-based protein expression in arugula protoplasts with 85% transformation efficiency. Lastly, we achieve 95% gene silencing in *Nb* leaves through CNT-mediated delivery of siRNA, whereby CNTs are shown to protect siRNA against nuclease degradation. This study establishes that CNTs, which are below the size exclusion limit of the plant cell wall (~ 20 nm), could be a promising solution for overcoming plant biomolecule delivery limitations in a species-independent and non-integrating manner and could enable high-throughput plant genetic transformations for a variety of plant biotechnology applications.

## Grafting DNA on carbon nanotube scaffolds

For the transgene expression study, we developed two distinct grafting methods to load green fluorescent protein (GFP)-encoding plasmids or their linear PCR fragments on SWCNTs and multi-walled carbon nanotubes (MWCNTs). The first DNA-grafting method involves direct adsorption of DNA on CNTs via dialysis. Initially, CNTs are coated with a surfactant – sodium dodecyl sulfate (SDS). During dialysis, SDS desorbs from the CNT surface and exits the dialysis membrane, while DNA adsorbs onto the surface of CNTs in a dynamic ligand exchange process (Fig. 1a). In this method, double-stranded DNA vectors graft on CNTs through π-π stacking interactions. The adsorption of DNA on CNTs is confirmed through a solvatochromic shift in the SWCNT nIR fluorescence emission spectra; characteristic of a DNA adsorption-induced change in the CNT dielectric environment^23^ (Supplementary Fig. 1). Control dialysis aliquots of SDS coated CNTs, in the absence of DNA, show rapid CNT precipitation and lack nIR fluorescence (Supplementary Fig. 1), confirming SDS desorption and replacement by DNA in our dialysis aliquots with DNA. Additionally, at the end of the dialysis procedure, we confirmed that there is no SDS left in the cartridge, by using Stains-all dye. Complete characterization (zeta potential, AFM height and DNA loading efficiency) of CNTs prepared via dialysis is summarized in Supplementary Fig. 1.

**Fig. 1.**
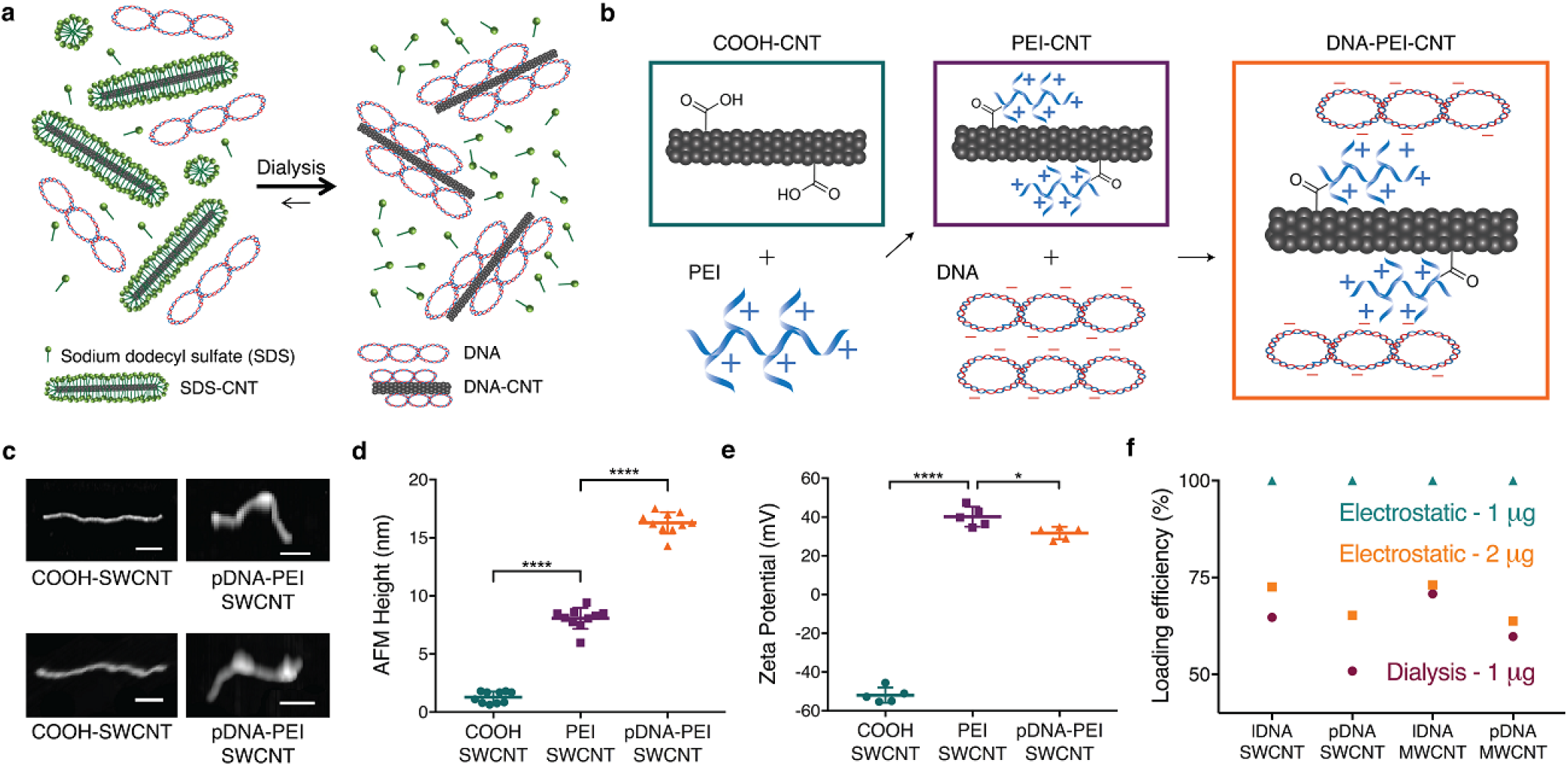
Strategies for grafting DNA on carbon nanotube scaffolds. **a**, DNA grafting on surfactant suspended CNTs through π-π stacking via dialysis method. **b**, DNA grafting on PEI modified carboxylated CNTs through electrostatic attachment. **c**, Representative AFM images of carboxylated SWCNTs and plasmid DNA loaded PEI modified SWCNTs. Scale bars, 100 nm. **d**, Average height profile of SWCNTs before and after PEI reaction and pDNA loading measured via AFM. \*\*\*\**P* < 0.0001 in one-way ANOVA. Error bars indicate s.d. (n = 10). e, Zeta potential measurements of SWCNTs before and after PEI reaction and pDNA loading measured via dynamic light scattering (DLS). \**P* = 0.0191 and \*\*\*\**P* < 0.0001 in one-way ANOVA. Error bars indicate s.d. (n = 5). f, Agarose gel electrophoresis results (Supplementary Fig. 1 and 2) demonstrate loading efficiency of 1 μg DNA onto 1 μg electrostatically modified and dialysis-made CNTs, and loading efficiency of 2 μg DNA onto 1 μg electrostatically modified CNTs.

The second method developed for DNA grafting on CNTs is electrostatic grafting, in which carboxylated CNTs (COOH-CNT) are first covalently modified with a cationic polymer (polyethylenimine, PEI) to carry a net positive charge. Next, positively charged CNTs (PEI-CNT) are incubated with negatively charged DNA vectors (Fig. 1b). The attachment of PEI and adsorption of DNA on CNTs is verified by atomic force microscopy (AFM) via CNT height increases (Fig. 1c). Nanoparticle heights before and after reaction with PEI are measured to be 1.3 nm and 8.1 nm for COOH- and PEI-SWCNT, respectively, confirming PEI binding. AFM also reveals that SWCNT height increases from 8.1 nm to 16.3 nm after incubation with DNA vectors, as expected, further confirming DNA grafting on SWCNTs (Fig. 1d). AFM characterization of MWCNT conjugates can be found in Supplementary Fig. 2.

The covalent attachment of PEI and electrostatic adsorption of DNA on CNTs is also confirmed through zeta potential measurements (Fig. 1e), after extensive washing of free unreacted PEI (see Methods). The initial zeta potential of −51.9 mV for COOH-SWCNT increases to +40.2 mV after reaction with positively-charged PEI, and subsequently decreases to +31.7 mV when incubated with negatively charged DNA, confirming PEI attachment and DNA adsorption. Complete characterization (zeta potential, AFM height and length, DNA loading efficiency) of electrostatically prepared CNT conjugates is summarized in Supplementary Fig. 2.

We note that DNA-CNT conjugates prepared via electrostatic grafting have higher DNA loading efficiencies compared to the conjugates prepared via the dialysis method. We demonstrate that the optimum DNA amount to be loaded on PEI-CNTs is with a 1:1 DNA:CNT mass ratio (Fig. 1f). Electrostatically grafted CNTs have 100% DNA loading efficiencies, whereas dialysis-loaded DNA-CNTs show 50-70% loading efficiencies when loaded with the same amount of DNA. Intracellular stability of DNA loaded PEI-SWCNT conjugates was tested by incubating conjugates with proteins (at total protein concentration similar to plant intracellular conditions), and even after 3 days of incubation, half of the DNA is found to remain adsorbed on the nanoparticles (Supplementary Fig. 2). We determined that PEI-CNTs are stable for 2 months at 4°C after their synthesis, and DNA loaded PEI-CNTs can be stored for a month at 4°C without compromising their stability. The 2-month stability of PEI-CNT complexes facilitates rapid loading of DNA onto CNTs through a 30-minute incubation of DNA vectors with PEI-CNTs.

## DNA delivery into mature plants with carbon nanotube scaffolds

Functional gene expression studies were implemented with arugula and cotton plant leaves to demonstrate the applicability of our platform to transform crop plants in addition to traditional model laboratory species, such as *Nicotiana benthamiana*. Furthermore, gene delivery and protein expression studies are carried out with wheat plants demonstrating that our platform is also applicable to transform monocot plant species in addition to dicot plants.

After preparation of DNA-CNT conjugates with GFP-encoding DNA plasmids or linear PCR amplicons with dialysis or electrostatic grafting, DNA-CNTs are infiltrated into the true leaves of mature plants (see Methods). Post-infiltration, DNA-CNTs traverse the plant cell wall and membrane to enter the cytosol. In the cytosol, we postulate that either the DNA-CNT complex further transports across the nuclear membrane where DNA desorbs from the CNT surface, or DNA desorbs from the CNT surface inside the cytosol and free DNA travels into the nucleus to initiate gene expression (Fig. 2a). Internalization of nanoparticles into mature plant cells is shown via tagging DNA-SWCNTs with a Cy3 fluorophore and tracking the Cy3-DNA-SWCNT conjugate inside the leaf tissue via confocal microscopy. For this experiment, a GFP mutant *Nb* plant that constitutively expresses GFP is used to co-localize the Cy3 fluorescence from the DNA-SWCNT with intracellular GFP fluorescence. When Cy3-DNA is delivered without SWCNTs, we do not observe overlay of Cy3 fluorescence with GFP, suggesting that Cy3-DNA is localized in the extracellular media. However, when Cy3-DNA-SWCNTs are delivered into the leaves, we observe 62% co-localization between the Cy3 and GFP fluorescence channels, which verifies efficient internalization of DNA-SWCNTs into plant cells (Fig. 2b). Internalization of nanoparticles into mature leaves is also shown with transmission electron microscopy (TEM) and nIR imaging of SWCNTs inside the leaf tissue (Supplementary Fig. 3).

**Fig. 2.**
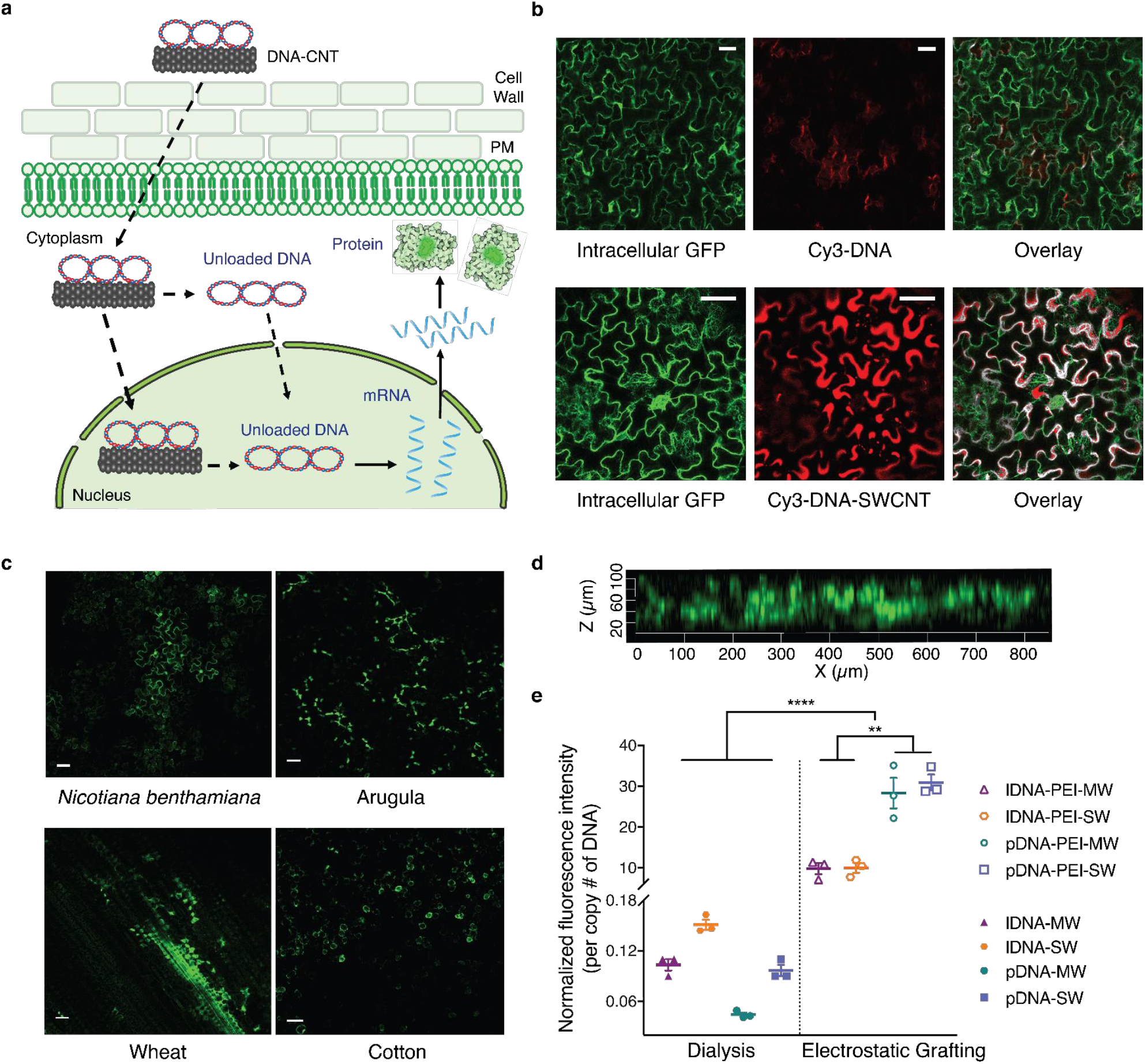
DNA delivery into mature plant leaves with CNTs and subsequent GFP expression. **a**, Schematic depicting DNA-CNT trafficking in plant cells and subsequent gene expression (dotted lines represent trafficking steps and the rigid lines represent gene expression steps). **b**, Nanoparticle internalization into mature plant cells is shown via imaging of Cy3 tagged DNA-SWCNTs with confocal microscopy, together with a control sample of Cy3 tagged DNA without SWCNTs. **c**, Wild type *Nb*, arugula, wheat, and cotton leaves infiltrated with DNA-CNTs are imaged with confocal microscopy for determining GFP expression levels in leaf lamina of each plant species. **d**, Z-stack analysis of the fluorescence profile of the DNA-CNT treated arugula leaf. **e**, Quantitative fluorescence intensity analysis of arugula confocal images for all nanomaterial formulations. \*\**P* = 0.001 and \*\*\*\**P* < 0.0001 in one-way ANOVA. Error bars indicate s.e.m. (n = 3). All scale bars, 50 μm.

Leaves infiltrated with DNA-CNTs for GFP expression were imaged with confocal microscopy, and expression of GFP is observed in the cells of the leaf lamina 72-hours post-infiltration in all plant species tested; *Nb*, arugula, wheat, and cotton (Fig. 2c). Z-stack analysis of the fluorescence profile of the DNA-CNT treated leaves shows that GFP fluorescence originates from the full thickness of the leaves, confirming that CNT nanocarriers diffuse and penetrate through the full leaf profile (Fig. 2d). No GFP expression is detected in the leaves when free DNA vectors, PEI-DNA complexes, or PEI-CNTs are delivered in control studies (Supplementary Fig. 4). Additionally, the spatial distribution of CNTs inside a leaf was modeled with a diffusion-reaction equation using the GFP expression profile as a proxy for nanocarrier diffusivity. The model predicts an exponential decay in the concentration of nanocarriers with respect to distance from the infiltration area, which agrees with the GFP expression profiles obtained via confocal imaging (Supplementary Fig. 5).

The efficiency of CNT nanocarrier internalization and GFP expression varies substantially for the different nanomaterial formulations we tested. Quantitative fluorescence intensity analysis of confocal images for arugula leaves (see Methods) indicates that GFP expression is significantly higher for DNA-CNTs prepared through electrostatic grafting compared to GFP expression induced by DNA-CNT conjugates prepared via π-π grafting with dialysis (Fig. 2e). Our most efficient DNA-CNT formulation is plasmid DNA delivered with PEI-functionalized SWCNT (pDNA-PEI-SWCNT), which is over 700 times more efficient than plasmid DNA adsorbed on pristine MWCNT via dialysis (pDNA-MWCNT), our least-efficient DNA-CNT formulation.

Our results suggest that the CNT surface chemistry is an important factor for biomolecule delivery into plant cells. The observed results can be explained by different DNA binding affinities to CNT surfaces in the two DNA grafting methods. The predominant DNA-CNT binding interaction in the case of dialysis is π-π stacking. In contrast, electrostatic attraction between PEI-CNTs and DNA is the predominant binding interaction for the electrostatic grafting method. We propose that the smaller equilibrium dissociation constant^24^ and higher binding energy value^25,26^ for electrostatic attraction compared to π-π stacking interactions increase the stability of the DNA-CNT complex as it traverses the cell wall, plasma membrane, and nuclear envelope, thus increasing the delivery efficiency of DNA to the plant cell. Worthy of note, for electrostatically-loaded DNA, we also observe statistically significantly higher protein expression efficiencies with CNT conjugates loaded with plasmid (pDNA) compared to linear (lDNA) conjugates. We hypothesize that this difference in transformation efficiency is due to the decreased cytosolic degradation rate of pDNA compared to lDNA in eukaryotic cells, as lDNA is prone to degradation by both endo- and exonucleases^27^, while pDNA is only degraded by endonucleases (as it does not contain free ends). Based on these results, all subsequent mature leaf transformation studies are performed with pDNA-PEI-SWCNTs, unless otherwise noted.

We further demonstrate that CNT-mediated gene expression is transient in mature plant leaves, independent of the plant species. Representative confocal images of pDNA-PEI-SWCNT infiltrated *Nb* (Fig. 3a) and corresponding quantitative fluorescence intensity analysis of these images demonstrate that the highest GFP fluorescence intensity at Day 3 disappears by Day 10 (Fig. 3b). Similarly, quantitative PCR (qPCR) analysis of GFP mRNA corroborates our confocal imaging results. For pDNA-PEI-SWCNT treated *Nb* leaves, we observe an over 7500-fold GFP mRNA increase 3-days post-infiltration, which drops to an insignificant two-fold mRNA change by Day 10 (Fig. 3c). Similar GFP expression profiles at Day 3 and 10 are also verified with arugula, wheat, and cotton mature leaves (Fig. 3d). Compared to CNT-mediated expression, however, *Agrobacterium-mediated* GFP expression in mature arugula leaves did not cease at Day 10, as shown by confocal imaging (Fig. 3e), GFP fluorescence intensity quantification (Fig. 3f), and qPCR analysis (Fig. 3g), supporting the established concept of DNA integration with *Agrobacterium*-mediated delivery^28^.

**Fig. 3.**
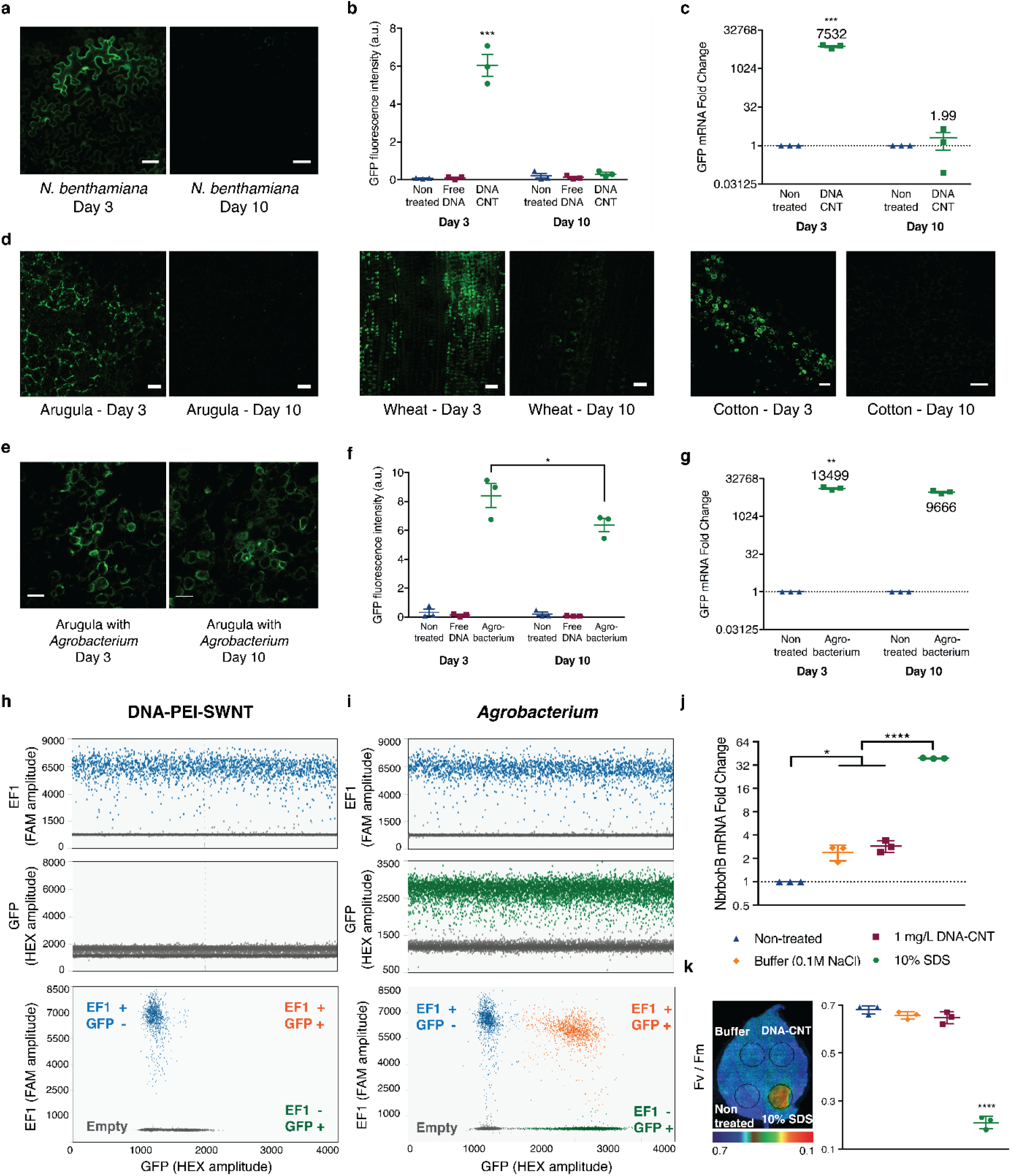
Transient CNT-mediated GFP expression in mature plant leaves and nanoparticle toxicity assessment. **a**, Representative confocal microscopy images of pDNA-PEI-SWCNT infiltrated mature *Nb* leaves imaged at Day 3 and 10. **b**, Quantitative fluorescence intensity analysis of confocal images at 3 and 10-days post-infiltration. \*\*\**P* = 0.0001 in two-way ANOVA. **c**, qPCR analysis of GFP mRNA expression levels at Day 3 and 10 in pDNA-PEI-SWCNT treated *Nb* leaves. \*\*\**P* = 0.0003 in two-way ANOVA. **d**, Representative confocal microscopy images at Day 3 and 10 in pDNA-PEI-SWCNT infiltrated mature arugula, wheat, and cotton leaves. **e**, Representative confocal microscopy images of *Agrobacterium* infiltrated mature *Nb* leaves imaged at Day 3 and 10. All scale bars, 50 μm. **f**, Quantitative fluorescence intensity analysis of *Agrobacterium*-transformed leaves at 3 and 10-days post-infiltration. \**P* = 0.012 in two-way ANOVA. **g**, qPCR analysis of *Agrobacterium*-transformed leaf at Day 3 and 10. \*\**P* = 0.0028 in two-way ANOVA. **h**, Droplet digital PCR (ddPCR) results of DNA-PEI-SWCNT infiltrated *Nb* leaves **i**, ddPCR results of *Agrobacterium* infiltrated *Nb* leaves. **j**, qPCR analysis of NbrbohB, a known stress gene in *Nb* plants, to test CNT toxicity. \**P* = 0.0169 and \*\*\*\**P* < 0.0001 in one-way ANOVA. **k**, Quantum yield measurements of photosystem II to test whether CNT-infiltrated leaves have similar photosynthesis quantum yield as control leaves without CNT infiltration. \*\*\*\**P* < 0.0001 in one-way ANOVA. All error bars indicate s.e.m. (n = 3).

Our results both at the mRNA transcript and fluorescent protein levels demonstrate that GFP expression is transient and suggest that genes delivered into plant cells via CNT nanocarriers do not integrate into the plant nuclear genome. We tested the non-integration of plasmid DNA into the plant nuclear genome via droplet digital PCR (ddPCR). ddPCR is a recently developed method that allows high-precision and absolute quantification of nucleic acid target sequences. One emerging application of ddPCR is to quantify rare genome editing events^29–33^, as ddPCR can accurately detect one copy of target DNA over 100,000 other DNA sequences, which makes ddPCR highly favorable tool to quantify low frequency gene integration events.

Here, we used ddPCR to determine whether DNA delivered with CNTs integrate into plant genomic DNA, and compared genomic DNA integration rates of CNT nanocarriers and *Agrobacterium*-mediated delivery methods. Our ddPCR experiments reveal that there is no transgene integration when DNA is delivered via CNTs (Fig. 3h), whereas high frequency GFP transgene integration events are shown when *Agrobacterium*-mediated delivery is performed (Fig. 3i). For each sample, the endogenous elongation factor 1 (EF1) gene is used as a reference gene (labeled with FAM), which is highly amplified both in DNA-PEI-SWCNT and *Agrobacterium* samples, whereas the target GFP gene (labeled with HEX) is only amplified in the *Agrobacterium* samples (Fig. 3h and i). We performed additional ddPCR control samples such as no template control (NTC), non-treated leaf, and free DNA infiltrated leaf. As expected, amplification of neither EF1 nor GFP genes is observed in the NTC, and amplification of only the EF1 gene is observed in non-treated or free DNA infiltrated leaves (Supplementary Fig. 6).

Biolistic (gene gun-based) DNA delivery is a preferred technique for transformation of plant species that are incompatible with *Agrobacterium*-based transformation. We next compared CNT-mediated DNA delivery with biolistic particle DNA delivery by transforming mature arugula true leaves and cotyledons with the same GFP-encoding plasmid using a gene gun. Interestingly, with biolistic transformation, we obtained little GFP expression in arugula true leaves, and also observed sparse GFP expression only in some of the guard cells on the topmost surface of arugula cotyledons through high-pressure condition biolistic delivery (Supplementary Fig. 7). Since GFP expression is limited to the topmost layer of the cotyledons, it is likely that biolistic delivery cannot penetrate deep enough in the arugula leaf to enable transformation of sub-cuticle cell types, such as mesophyll cells, even though a wide range of gene gun pressures (up to 900 PSI) were also tested. To our knowledge, biolistic transformation of arugula leaves remains to be shown in the literature. Consequently, we tested the transformation of mature *Nb* plant leaves with biolistic transformation and successfully obtained GFP expression in mesophyll cells, most likely due to the fact that, as a model laboratory plant, *Nb* has a thin and easy-to-penetrate leaf structure^34^ (Supplementary Fig. 7). These results demonstrate that depending on the plant species and tissue type, biolistic transformation can result in variable tissue penetration depth and expression efficiency, unlike CNT-mediated delivery.

## Testing CNT toxicity or tissue damage in plant leaves

To test whether CNT nanocarriers are biocompatible, we undertook plant toxicity and tissue damage tests. Specifically, for toxicity analyses, we performed qPCR analysis of respiratory burst oxidase homolog B (NbrbohB) upregulation (Fig. 3j), a known stress gene representing many different types of stress conditions (mechanical, light, heat, biotic, etc.) in *Nb* plants^35^. Quantification of NbrbohB expression shows that CNT-treated areas in leaves do not upregulate NbrbohB compared to adjacent areas within the same leaves treated only with buffer. Additionally, quantum yield measurements of photosystem II^36^ show that CNT-infiltrated areas in *Nb* leaves have similar photosynthesis quantum yields as control areas within the same leaves that were infiltrated only with buffer, or not infiltrated at all (Fig. 3k). Positive controls to induce plant stress for both NbrbohB qPCR and photosystem II quantum yield measurements show clear upregulation of NbrbohB and a significant decrease in photosystem II quantum yield in *Nb*. We also analyzed leaf tissue damage visually and via confocal microscopy, which again show no sign of tissue damage in CNT-infiltrated leaves (Supplementary Fig. 8). Our analysis concludes that the CNT-based delivery platform is biocompatible and does not induce toxicity or tissue damage to mature plants with the conditions used in the present study.

## DNA delivery into isolated protoplasts with carbon nanotube scaffolds

We further investigated the ability of CNT nanocarriers to deliver plasmid DNA and trigger functional gene expression in a different plant system – isolated protoplasts, which are cultured plant cells without cell walls. Currently, protoplasts are used to increase the throughput of plant genetic screens and for synthesis of recombinant proteins, thus benefiting from a facile, passive, high efficiency, and species-independent transformation platform^37^. For this purpose, intact and healthy protoplasts were extracted from arugula leaves through enzymatic cell wall degradation (Fig. 4a) with high efficiency and high yield (10^7^ total protoplasts / 10 leaves). Nanoparticle internalization into isolated protoplasts was confirmed via nIR imaging of DNA-SWCNTs by taking advantage of the intrinsic nIR fluorescence of SWCNTs that emits in a separate optical window from chlorophyll broadband autofluorescence. When DNA-SWCNTs are added to a protoplast solution, we observe nIR SWCNT fluorescence that colocalizes with the brightfield image of the protoplast, confirming SWCNT internalization. Conversely, without DNA-SWCNT addition, no nIR fluorescence is observed (Fig. 4b).

**Fig. 4.**
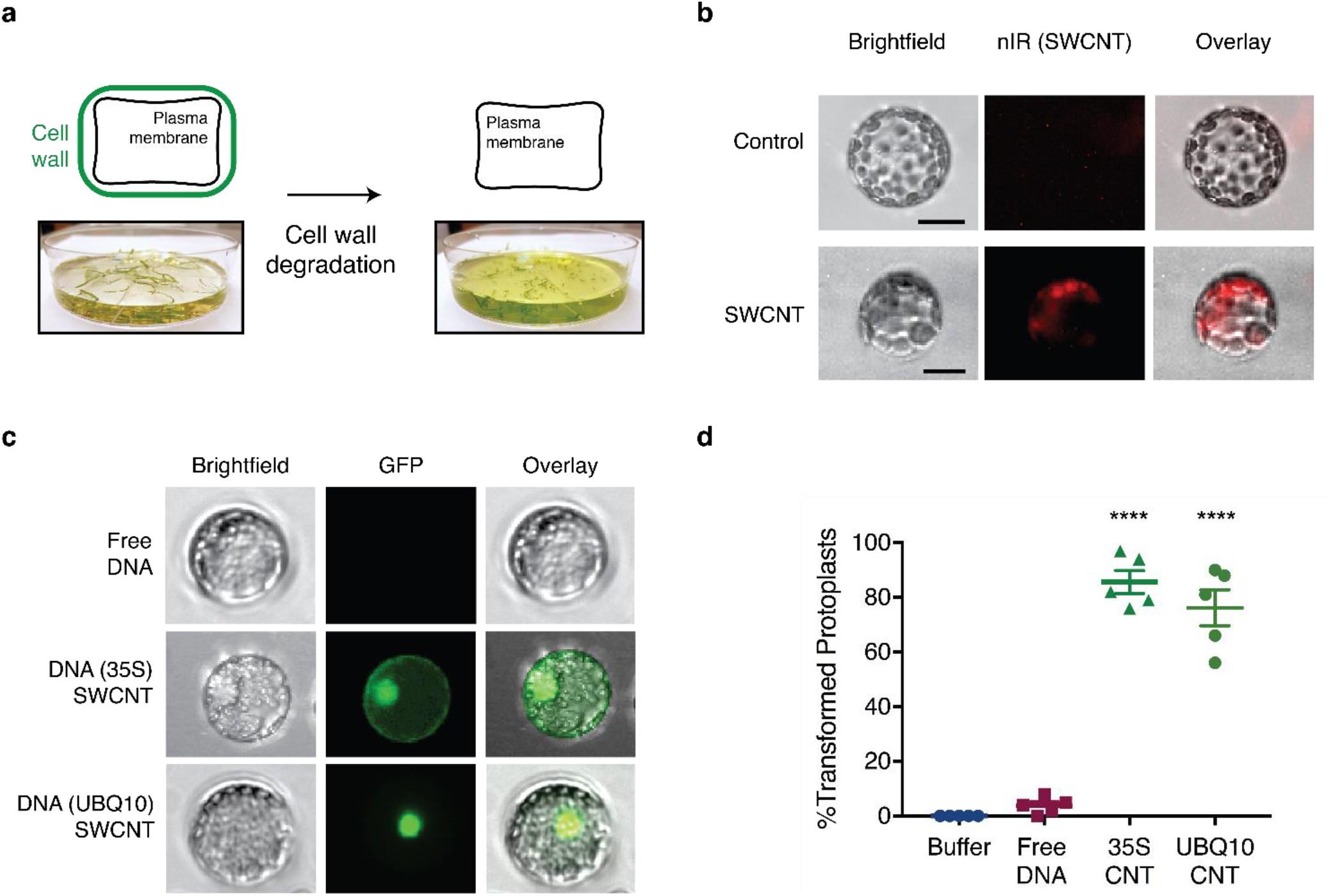
DNA delivery into isolated protoplasts with CNTs and subsequent GFP expression. **a**, Intact and healthy protoplast extraction from arugula leaves through enzymatic cell wall degradation. **b**, Nanoparticle internalization into isolated protoplast verification by imaging the intrinsic nIR fluorescence of SWCNTs after incubation with protoplasts. Scale bars, 10 μm. **c**, GFP expression imaging of protoplasts incubated with 35S and UBQ10 plasmid carrying DNA-CNTs via fluorescence microscopy. Protoplast diameters are ~20 μm. **d**, Percentage of the total isolated protoplasts transformed with 35S-CNTs and UBQ10-CNT after a 24-hour incubation period. **** *P* < 0.0001 in one-way ANOVA. Error bars indicate s.e.m. (n = 5).

For gene expression studies, isolated protoplasts were incubated with plasmid DNA-SWCNTs prepared via dialysis, and subsequently imaged with fluorescence microscopy. In addition to the plasmid used in leaf studies (35S-GFP), for protoplast experiments we also used a plasmid that encodes a nuclear localization signal (UBQ10-GFP, Supplementary Fig. 9), which transports the expressed GFP protein from the cytosol into the nucleus. Protoplasts incubated with both types of DNA-CNTs show GFP expression correctly localized in cells, whereas protoplasts incubated with free plasmids without CNTs do not show GFP expression (Fig. 4c). CNT-mediated protoplast transformation efficiencies are 76% and 86% with UBQ10-CNTs and 35S-CNTs, respectively (Fig. 4d and Supplementary Fig. 10). Our prior work on CNT internalization into extracted chloroplasts suggests nanoparticle internalization through the lipid bilayer occurs within seconds of CNT exposure^15^. Thus, our CNT-based plasmid DNA delivery platform enables rapid and passive delivery of DNA into protoplasts and transgene expression with high efficiency and no observable adverse effects to protoplast viability (Supplementary Fig. 10).

## Carbon nanotube-guided siRNA gene silencing in mature plants

We next demonstrate the applicability of our CNT-mediated delivery tool in plants with another broadly-utilized functional cargo – siRNA, which is a short RNA duplex that acts within the RNA interference (RNAi) pathway for sequence-specific inhibition of gene expression at the mRNA transcript level^38^. As with plasmid DNA, delivery of siRNA has been optimized for most mammalian and bacterial cell culture applications, but many aspects of siRNA delivery to mature plants still remains a significant challenge^39^.

For this study, we silence a gene in transgenic *Nb*, which strongly expresses GFP in all cells due to GFP transgene integration in the *Nb* nuclear genome. To silence this constitutively-expressed GFP gene, we designed a 21-bp siRNA sequence that is specific to the GFP mRNA^40^ (Fig. 5a). Loading of siRNA on CNTs is accomplished by probe-tip sonication of each siRNA single-strand (sense and antisense) with SWCNTs (see Methods and Fig. 5a). The adsorption of RNA on SWCNTs was confirmed through the emergence of characteristic peaks in the SWCNT nIR fluorescence emission spectra for both siRNA sense and antisense suspensions (Supplementary Fig. 11). The mixture of siRNA sense and antisense loaded CNTs was infiltrated into the leaves of mature transgenic *Nb* plants. Post-infiltration, we predict that RNA-CNTs traverse the plant cell wall and membrane, and reach the cytosol. In the plant cell cytosol, the complementary siRNA strands hybridize to each other, desorb from the CNT surface, and induce GFP gene silencing through the RNAi pathway (Fig. 5a). Cytosolic hybridization and desorption claims are supported by our thermodynamics analysis that considers the energetics of hydrogen bonding and π-π stacking interactions, which found out to favor siRNA hybridization and desorption over adsorption on CNTs only in the intracellular environment (see Supplementary Information and Supplementary Fig. 11).

**Fig. 5.**
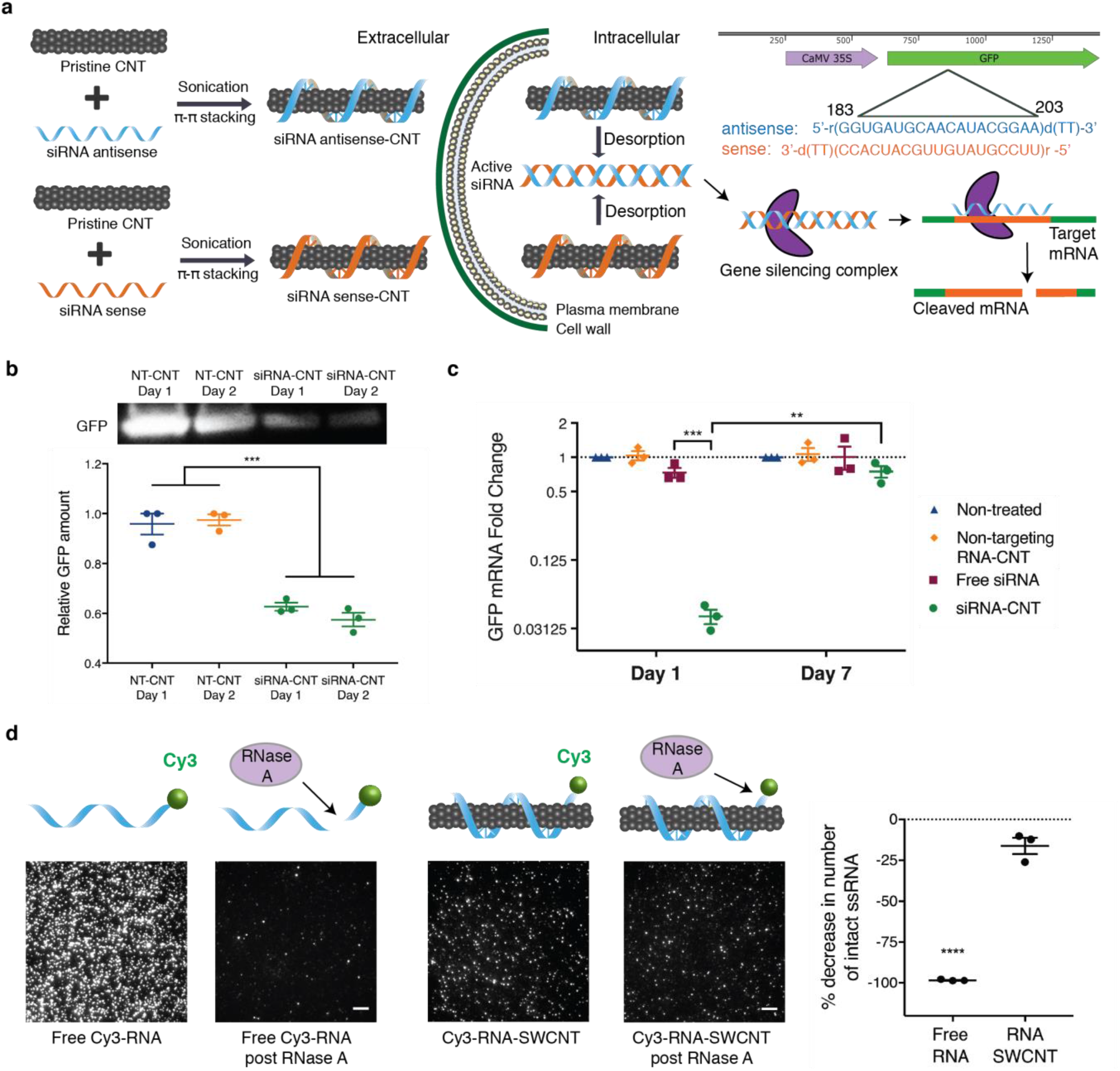
CNT-guided siRNA gene silencing in mature plants. **a**, Loading of siRNA on CNTs through probe-tip sonication of sense and antisense sequences with CNTs. Schematic depicting RNA-CNTs trafficking in cells and subsequent gene silencing through RNAi pathway. **b**, A representative Western blot showing GFP extracted from non-targeting RNA-CNTs (NT-CNT) and siRNA-CNT treated leaves at one and two days post-infiltration. \*\*\**P* = 0.0001 in one-way ANOVA. **c**, qPCR results for non-treated leaf and leaves infiltrated with NT-CNT, free GFP-targeting siRNA, and GFP-targeting siRNA delivered via CNT scaffolding at one and seven day post-infiltration. \*\**P* = 0.0012 and \*\*\**P* = 0.0002 in two-way ANOVA. All error bars indicate s.e.m. (n = 3). **d**, Single molecule TIRF microscopy of Cy3-labeled RNA and Cy3-labeled RNA-SWCNTs before and after incubation with RNase A. Scale bars, 5 μm. Fluorophore counts for each sample were normalized with respect to the counts in a channel flushed with salt solution to account for washing away of surface immobilized particles by lateral flow. All error bars indicate s.e.m. (n = 3). \*\*\*\**P* < 0.0001 in two-tailed unpaired t-test.

Transgenic *Nb* leaves that constitutively express GFP were imaged via confocal microscopy to monitor GFP silencing at the protein level. Untreated leaves show strong GFP expression, as expected, due to the constitutive expression of GFP in the transgenic plant. Conversely, leaves infiltrated with siRNA-CNTs show reduced GFP fluorescence via confocal microscopy (Supplementary Fig. 12). Moreover, Western blot analysis reveals 43% reduction in GFP protein extracted from siRNA-CNT treated leaves compared to the leaves treated with non-targeting RNA loaded CNTs (NT-CNT) at two days post-infiltration (Fig. 5b). To corroborate confocal imaging and Western blot results, we performed qPCR analysis of the siRNA-CNT infiltrated plant leaf tissue to quantify silencing at the mRNA transcript level. No significant GFP mRNA reduction is observed in the non-treated leaf, nor in leaves infiltrated with non-targeting RNA-CNTs, nor in leaves infiltrated with free GFP-targeting siRNA (Fig. 5c), whereby 95% reduction in GFP mRNA is observed at Day 1 when GFP-targeting siRNA is delivered via CNT scaffolding. 7-days following the introduction of siRNA-CNTs, GFP expression as measured by qPCR returns to baseline levels as observed in non-treated leaves (Fig. 5c). A separate qPCR trial shows we are able to recover 71% GFP silencing at Day 7 through the re-infiltration of the siRNA-CNT suspension at Day 5 (Supplementary Fig. 12).

It is likely that CNT scaffolding improves internalization of siRNA and also protects siRNA from degradation once intracellular. To explore this hypothesis, we performed single molecule total internal reflection fluorescence (smTIRF) microscopy to probe single siRNA susceptibility to degradation by RNase A when adsorbed on SWCNTs, compared to free siRNA. To do so, we labeled the antisense strand of GFP siRNA with a Cy3 fluorophore, and immobilized RNA-Cy3 and RNA-Cy3-SWCNTs onto parallel microfluidic channels of a microscope slide (see Methods). We measured the Cy3 fluorescence in each channel before and after treatment with RNase A, whereby percent fluorescence decrease was used as a proxy for the percent siRNA degraded. Our TIRF results show that 98% of the initial Cy3-RNA immobilized on the channel surface is degraded after incubation with RNase A, whereas only 16% of Cy3-RNA is degraded when it is bound to SWCNTs, suggesting that CNTs can protect the siRNA cargo from metabolic degradation inside cells (Fig. 5d). Negative controls in which only salt buffer is flown through, or empty BSA-passivated channels, do not show appreciable changes in fluorescence or fluorescence counts, respectively (Supplementary Fig. 13).

## Conclusions

Genetic engineering of plants may address the crucial challenge of cultivating sufficient food, natural product therapeutics, and bioenergy for an increasing global population living under changing climatic conditions. Despite advances in genetic engineering across many biological species, the transport of biomolecules into plant cells remains as one of the major limitations for rapid, broad-scale and high-throughput implementation of plant genetic engineering, particularly for intact plant tissues and organs. We thus present a nanomaterial-based delivery platform that permits diverse conjugation chemistries to achieve DNA delivery without transgene integration in both model and crop plants, and in both dicot and monocot plants, with high efficiency and without toxicity or tissue damage. In this study, we show the development and optimization of dialysis and electrostatic grafting methods for loading biomolecules onto high aspect ratio CNTs. We confirm the feasibility and test the efficacy of this platform by delivering reporter GFP DNA constructs into mature *Nicotiana benthamiana*, arugula, wheat, and cotton leaves, and arugula protoplasts, and obtain strong expression of a functional transgenic protein. The value of the developed platform is also demonstrated through high efficiency gene silencing obtained via CNT-mediated siRNA delivery in mature *Nb* leaves.

The nanomaterial-based transient plant transformation approach demonstrated herein is beneficial for plant biotechnology applications where gene expression without transgene integration is desired, and is amenable to multiplexing whereby multiple gene vectors are to be delivered and tested rapidly in a combinatorial manner and in parallel^41^. This approach may aid high-throughput screening in mature plants to rapidly identify genotypes that result in desired phenotypes, mapping and optimization of plant biosynthetic pathways, and maximization of plant-mediated natural product synthesis, most of which currently rely on *Agrobacterium*-mediated transformation^42^. CNT-mediated delivery is well-suited to transient testing of plant transgenes for such applications as it is easy, cost effective, non-destructive, fast, species-independent, and scalable.

Additionally, global regulatory oversight for genetically-modified organisms (GMOs) motivate the future development of non-integrative and/or DNA-free plant genetic transformation approaches in which the delivered gene expression is transient and foreign DNA is not integrated into the plant genome^43^. However, the most commonly used tool today for plant genetic transformations – *Agrobacterium*-mediated transformation technology – is unable to perform DNA- and transgene-free editing, yields random DNA integration, and has highly limited plant host range. If combined with nuclease-based genome editing cargoes such as ZFNs, TALENs, Cpf1, and CRISPR/Cas9, CNTs can enable transient expression of these tools for production of heritable edits. As such, CNT-based delivery of these biomolecular cargoes could enable high-efficiency genome modification without transgene integration, the latter of which results from either *Agrobacterium* or biolistic-mediated plant genetic transformation, thus falling under the purview of GMO regulations. This latter application of the presented technology will be particularly beneficial for heterogeneous plant species such as cassava, cacao, sugarcane, etc. in which crossing cannot be used to remove transgenes. Furthermore, CNTs are shown herein to protect RNA cargo against nuclease degradation, a feature of CNT-based delivery that may be extended to the protection of other biological cargoes of interest.

Thus, in this study, we develop plant transformation biotechnologies that show high efficiency, species-independence, and which can be used as a transgene-free plant genetic engineering approach to keep pace with a growing need for efficient delivery of biomolecules to plants that are not subject to regulatory oversight as genetically modified organisms.

## Acknowledgments

We acknowledge support of a Burroughs Wellcome Fund Career Award at the Scientific Interface (CASI), a Stanley Fahn PDF Junior Faculty Grant with Award # PF-JFA-1760, a Beckman Foundation Young Investigator Award, a USDA AFRI award, and a FFAR New Innovator Award (M.P.L). M.P.L. is a Chan Zuckerberg Biohub investigator. G.S.D. is supported by a Schlumberger Foundation Faculty for the Future Fellowship. L.C. is supported by National Defense Science and Engineering Graduate (NDSEG) Fellowship and by the LAM Foundation. The authors wish to thank Christopher Gee for assisting with the Imaging-PAM Maxi fluorimeter, Dr. Alex Schultink and Arturo Ortega for helpful discussions, and Dr. Christopher Jakobson and Dr. Danielle Tullman-Ercek for generously sharing their lab resources. We acknowledge the support of Dr. Holly Aaron at the UC Berkeley Molecular Imaging Center, the QB3 Shared Stem Cell Facility, and the Innovative Genomics Institute (IGI).

## Author contributions

G.S.D. and M.P.L. conceived of the project, designed the study, and wrote the manuscript. G.S.D. performed the majority of experiments and all data analysis. H.Z. and L.C. performed AFM imaging, and H.Z. also performed nanoparticle internalization experiments into mature leaves and Western blot experiments. J.L.M. performed *Agrobacterium* and wheat transformation experiments. N.G. helped designing ddPCR experiments, performed CNT leaf toxicity confocal imaging and TIRF experiments. F.C. performed nanoparticle internalization experiments into isolated protoplasts. Y.S. performed TEM imaging of leaves. A.A. prepared plasmids used in the studies, and R.C. prepared siRNA-SWCNTs conjugates. M.-J.C. performed particle bombardment experiments. All authors have edited and commented on the manuscript, and have given their approval of the final version.

## Competing interests

The authors declare no competing interests.

**Correspondence and requests for materials** should be addressed to M.P.L.

## Methods

### Procurement and preparation of chemicals and nanomaterials

Super purified HiPCO SWCNTs (Lot # HS28-037) were purchased from NanoIntegris, MWCNTs (Lot # R0112) were purchased from NanoLab, and both CNT samples were extensively purified before use^1^. Carboxylic acid functionalized SWCNTs (Lot # MKBX0303V) and MWCNTs (Lot # BCBR9248V) were purchased from Sigma-Aldrich. GFP-encoding dicot plasmids (35S-GFP-NOS and UBQ10-GFP-NOS) were obtained from the Sheen Lab, Harvard Medical School^2^. GFP-encoding monocot plasmid (osACTIN-GFP-NOS) was obtained from the Staskawicz Lab, UC Berkeley. 20K MWCO dialysis cassettes were purchased from Thermo Scientific. The following chemicals were purchased from Sigma-Aldrich: stains-all dye (95%), sodium dodecyl sulfate (molecular biology grade), sodium chloride, MES hydrate, D-mannitol, calcium chloride dihydrate (suitable for plant cell culture), potassium chloride, magnesium chloride hexahydrate, bovine serum albumin (heat shock fraction), polyethylene glycol (4 K), and polyethylenimine (branched, 25 K). Cellulase R10 and macerozyme R10 enzymes were purchased from Grainger. Single stranded RNA and DNA polymers were purchased from IDT and dissolved in 0.1M NaCl before use. All ddPCR reagents and materials were purchased from Bio-Rad. BSA-Biotin and NeutrAvidin were purchased from Sigma-Aldrich and RNase A was purchased from Takara Bio. UltraPure DNase/RNase-free distilled water from Invitrogen was used for qPCR and ddPCR experiments, and EMD Millipore Milli-Q water was used for all other experiments.

### Plant growth

Italian arugula (*Eruca sativa*) seeds purchased from Renee’s Garden were germinated in SunGro Sunshine LC1 Grower soil mix by planting the seeds half an inch deep into the soil of a standard propagation liner tray (Nursery Supplies). The germinated plants were then moved to a Hydrofarm LED growth chamber (12h light at ~22°C / 12h dark at 18°C). Plants were allowed to mature to 3-4 weeks of age within the chamber before experimental use. Wild type and transgenic mGFP5 *Nb* seeds obtained from the Staskawicz Lab, UC Berkeley, were germinated and grown in SunGro Sunshine LC1 Grower soil mix for four weeks before experimental use. Spring wheat (*Triticum aestivum* L., cv. Fielder) were grown in Supersoil (Rod McClellan Co., South San Francisco, CA, USA) in a Conviron growth chamber with 60% relative humidity, 18-hour light at 24°C: 8-hour dark at 18°C cycle, and 3-4-week-old plants were used for experiments. Cotton seedlings were purchased from Cottonman.com and allowed to mature within the Hydrofarm LED growth chamber (12h light at ~22°C / 12h dark at 18°C).

### SDS-CNT, siRNA-CNT and ssDNA-CNT preparation

3 mg HiPCO SWCNTs were added to 3 mL 2 wt% SDS in water and bath sonicated for 10 min, followed by probe-tip sonication with a 6-mm sonicator tip at 10% amplitude for 30 min in an ice bath (pulse 1 sec on /1 sec off). The resulting solution rested at room temperature for 30 minutes before centrifugation at 16,100g for 1 h to remove unsuspended SWCNT aggregates and metal catalyst precursor. The concentration of SDS-SWCNTs (supernatant) was measured by recording the SWCNT absorption spectrum with a UV-Vis-nIR spectrometer and calculating the SWCNT concentration in mg/liter (absorbance at 632 nm/extinction coefficient of 0.036). The same suspension protocol applies for MWCNTs, but, their concentration was measured using a standard curve as obtained by Yang *et al*. ^3^.

The sequences of siRNA that were utilized for siRNA gene silencing experiments are: sense strand: 5’-r(GGUGAUGCAACAUACGGAA)d(TT)-3’ and antisense strand: 5’-r(UUCCGUAUGUUGCAUCACC)d(TT)-3’.

The sequences of the non-targeting RNA strands are sense: 5’-r(UAAGGCUAUGAAGAGAUAC)d(TT)-3’ and antisense: 5’-r(GUAUCUCUUCAUAGCCUUA)d(TT)-3’.

siRNA and non-targeting RNA were loaded on SWCNTs as single-stranded polymers through probe-tip sonication as previously described^4^. Briefly, the sense strand of siRNA was dissolved at a concentration of 100 mg/mL in 0.1 M NaCl. 20 μL of this RNA solution was aliquoted into 980 μL 0.1 M NaCl and 1 mg HiPCO SWCNTs was added. The mixture was bath sonicated for 10 min, followed by probe-tip sonication with a 3-mm tip at 50% amplitude (~7W) for 30 min in an ice bath. The resulting solution rested at room temperature for 30 minutes before centrifugation at 16,100g for 1 h to remove unsuspended SWCNT aggregates and metal catalyst precursor. The same protocol was followed for the antisense strand of siRNA and non-targeting RNA strands. Unbound (free) RNA was removed via spin-filtering (Amicon, 100 K) 10-15 times and the concentration of RNA-SWCNTs was determined by measuring the SWCNT absorbance at 632 nm. For toxicity, tissue damage and Cy3-DNA internalization assays, SWCNTs were suspended in single-stranded DNA (ssDNA) polymers with the (GT)15 sequences. Preparation of ssDNA-SWCNTs followed the same protocol as for RNA-loaded SWCNTs, described above.

### Linear DNA vector preparation from plasmid DNA

The promoter, GFP gene, and terminator regions of the 35S-GFP-NOS plasmid (obtained from the Sheen Lab, Harvard Medical School) were amplified with PCR over 35 cycles, with the following modified M13 forward and M13 reverse primers: 5′-GTAAAACGACGGCCAGT-3′ and 5′-AGCGGATAACAATTTCACACAGG-3′, respectively. Following PCR, pure DNA vector was obtained by using a PureLink PCR purification kit (Invitrogen) to eliminate primers, unreacted nucleotides, and enzymes. To check the amplification quality, the resulting amplicon was sent for Sanger sequencing, and was also run with agarose gel electrophoresis (see Supplementary Fig. 7 for plasmid maps and linearization results).

### Direct adsorption of DNA onto CNTs via dialysis

SDS-CNT solution containing 1 μg of CNTs, and 10 μg of free plasmid DNA were placed into an accurately-pore-sized dialysis cartridge (20 kDa MWCO, 0.5 mL), that allowed the exit of SDS monomers that desorbed from the CNT surface, while free plasmid DNA suspended the CNTs which remained inside the dialysis cartridge. If necessary due to volume considerations, 2 wt% SDS was used to fill the additional volume of dialysis cartridge to ensure there was no free air space in the cartridge. After 4 days of dialysis with continuous stirring at room temperature and changing the dialysis buffer (0.1M NaCl) daily, we obtained a stable suspension of plasmid DNA conjugated CNTs. The preparation protocol was same for both plasmids and linearized DNA vectors, and for both types of CNTs (SW- and MWCNTs). The nIR fluorescence spectra of dialysis suspended CNTs are obtained through a nIR fluorescence microscopy using 721nm laser excitation and an inverted microscope outfitted with an InGaAs sensor array for imaging^4^.

#### Control studies for dialysis

A control cartridge consisting of an SDS-CNT solution that contains 1 μg of CNTs in 2 wt% SDS, but lacking DNA, was dialyzed in parallel under the same conditions to ensure that CNTs did not suspend in solution in the absence of plasmid DNA, confirming plasmid DNA adsorption to CNTs in the main sample. Stains-all dye, which changes color in the presence of SDS, is used to determine %SDS in the dialysis cartridge. A standard curve with the range of 0-0.016 %SDS is created at the absorbance wavelength 453 nm. Five dialysis formulations, as described above, are prepared and they are stopped at different time points along the duration of dialysis (Day 0, 1, 2, 3 and 4). 10 μL of dialysis solution is mixed with 1 mL 0.1% stains-all (w:v in formamide), and absorbance at 453 nm is measured. By using the standard curve, precise SDS% in the cartridge was calculated at each day point.

### Electrostatic grafting of DNA onto CNTs

Chemical modification of CNTs to carry positive charge is described elsewhere^5^ and applied here with some modifications. 10 mg COOH-CNT powder was added to 10 ml water (can be scaled up or down as desired at 1mg/ml concentration). The solution was bath sonicated for 5 minutes and probe-tip sonicated with 6-mm tip at 10% amplitude for 30 min on ice. It was rested 30 minutes at room temperature and centrifuged at 16,000g for 1 h. Supernatant was taken and SWCNT concentration was measured via absorbance at 632 nm with extinction coefficient of 0.036 to convert to mg/L. MWCNT concentration was measured using a standard curve as obtained by Yang *et al*.^3^. Prepared COOH-CNT solution was mixed with PEI at a mass ratio of 1:10 CNT:PEI. The solution was bath sonicated for several minutes, and subsequently heated at 82 °C with stirring for 16 h (the reaction can be scaled up or down as desired by keeping the PEI to CNT mass ratio constant). The reaction mixture was subsequently cooled to room temperature and filtered with a 0.4 μm and 1 μm Whatman Nucleopore membrane to filter SWCNTs and MWCNTs, respectively. The filtered product was washed vigorously with water 10 times to remove unreacted PEI from the reaction mixture, then dried and collected. 3 mg of dried product (PEI-CNT) was subsequently suspended in 3 mL water by probe-tip sonication with a 6-mm tip at 10% amplitude for 30 min in an ice bath. The resulting solution was rested at room temperature for 30 minutes before centrifugation at 16,100g for 1 h to remove unsuspended CNT aggregates. The PEI-CNT solution containing 1 μg of CNTs was added into 1 μg of DNA dropwise, pipetted in and out 10 times, and incubated at room temperature for 30 minutes (DNA incubation can be scaled up or down by keeping the DNA to PEI-CNT mass ratio constant).

### AFM characterization

3 μL of sample was deposited on a freshly cleaved mica surface and left to adsorb on the surface for 5 minutes. The mica surface was then slowly rinsed with water for three times (each time with 10 μL water) to remove the salt. Then, the mica surface was dried with a mild air stream by an ear-washing bulb and was imaged with a MultiMode 8 AFM with NanoScope V Controller (Bruker, Inc.) under tapping mode in air. All AFM images were analyzed by NanoScope Analysis v1.50.

### Infiltration of leaves with CNTs

Healthy and fully-developed leaves from arugula (3-4 week old), *Nicotiana benthamiana* (4 week old), wheat (4 week old), and cotton (4 week old) plants were selected for experiments. A small puncture on the abaxial surface of the arugula and cotton leaf lamina was introduced with a pipette tip, and 100-200 μL of the plasmid DNA-CNT solution (or of any control solution) was infiltrated from the hole with a 1 mL needleless syringe by applying a gentle pressure, with caution not to damage the leaf. For *Nb* infiltration, a tiny puncture on the abaxial surface of the leaf lamina was introduced with a sharp razor, and 100-200 μL of siRNA-CNT or DNA-CNT solution (or of any control solution) was infiltrated through the puncture with a 1 mL needleless syringe by applying a gentle pressure. After infiltration, leaves were left in the plant growth chamber (for arugula and cotton) and in plant pots (for other plants) to allow for gene expression and silencing, and imaged after 72 and 24 h, respectively, prior to quantifying gene expression and silencing. For wheat leaf infiltrations, a sharp razor blade was used to produce a small puncture on the abaxial surface of 3-4-week-old plant leaves, and 100-200 μL of the plasmid DNA-CNT solution (or of any control solution) was infiltrated with a 1 mL needless syringe. Plants were returned to growth chamber and imaged after 3 and 10 days-post-infiltration.

### Quantitative fluorescence intensity analysis of GFP gene expression

DNA-CNT Infiltrated plant leaves were prepared for confocal imaging 72-hours post-infiltration by cutting a small leaf section of the infiltrated leaf tissue, and inserting the tissue section between a glass slide and cover slip of #1 thickness. 100 μL water was added between the glass slide and cover slip to keep the leaves hydrated during imaging. A Zeiss LSM 710 confocal microscope was used to image the plant tissue with 488 nm laser excitation and with a GFP filter cube. GFP gene expression images were obtained at 10x magnification. Confocal image data was analyzed to quantify GFP expression across samples. For each sample, 3 biological replicates (3 infiltrations into 3 different plants) were performed, and for each biological replicate, 15 technical replicates (15 non-overlapping confocal field of views from each leaf) were collected. Each field of view was analyzed with custom ImageJ analysis to quantify the GFP fluorescence intensity value for that field of view, and all 15 field of views were then averaged to obtain a mean fluorescence intensity value for that sample. The same protocol was repeated for all 3 biological replicates per sample, and averaged again for a final fluorescence intensity value, which correlates with the GFP expression produced by that sample.

### Quantitative PCR (qPCR) experiments for gene expression

Two-step qPCR was performed to quantify GFP gene expression in wild type *Nb* plants with the following commercially-available kits: RNeasy plant mini kit (QIAGEN) for total RNA extraction from leaves, iScript cDNA synthesis kit (Bio-Rad) to reverse transcribe total RNA into cDNA, and PowerUp SYBR green master mix (Applied Biosystems) for qPCR. The target gene in our qPCR was GFP, and the reference gene was elongation factor 1 (EF1). Primers for these genes were ordered from IDT. GFP primers used are: forward 5’-CGCCGAGGTGAAGTT-3’ and reverse: 5’-GTGGCTGTTGTAGTTGTAC-3’. Primers for EF1 are: forward: 5’-TGGTGTCCTCAAGCCTGGTATGGTTGT-3’ and reverse: 5’-ACGCTTGAGATCCTTAACCGCAACATTCTT-3’. An annealing temperature of 60°C was used for qPCR, which we ran for 40 cycles. qPCR data was analyzed by the ddCt method^6^ to obtain the normalized GFP gene expression-fold change with respect to the EF1 housekeeping gene and control sample. For each sample, qPCR was performed as 3 technical replicates (3 reactions from the same isolated RNA batch), and the entire experiment consisting of independent infiltrations and RNA extractions from different plants was repeated 3 times (3 biological replicates).

### *Agrobacterium-mediated* transformation

*Agrobacterium tumefaciens* strain GV3101 was used for genetic transformation of *Nb* and arugula leaves, and as a positive control in ddPCR experiments. To generate the *Agrobacterium*-binary construct, the DNA fragment containing 35S-GFP-NOS were excised from the plasmid 35sC4PPDKsGFPTYG with the restriction enzymes XhoI and EcoRI and cloned into an entry cloned digested with the same restriction enzymes. The 35S-GFP-NOS entry clone was recombined into the *Agrobacterium* destination vector pPZP201^7^. Agrobacterium suspensions (OD600 = 0.4) were infiltrated into *Nb* and arugula leaves of 3-4-week-old plants using a 1-ml needleless syringe. Plants were returned to the growth chamber and imaged after 3 and 10 days-post-infiltration, and used in ddPCR experiments 14-days post-infiltration.

### Biolistic delivery of plasmid DNA

*Nb* and arugula seeds were sterilized in solution (20% bleach) for 30 minutes under gentle agitation, then washed three times with sterile water, plated on ½ strength Murashige and Skoog (MS) medium, stratified for 2 days at 4°C before transferring to a 26°C incubator with 16-hour light: 8-hour dark cycle for growth. 3-wk-old leaves were placed onto semi-solid pre-shooting media [4.43 g/L of MS basal medium and vitamins; 36.43 g/L of mannitol; 36.43 g/L of sorbitol; 0.30g of casein enzymatic hydrolysate; 0.5 g/L of L-proline 2 mL/L of 2,4-D (1 mg/ml); pH 5.8; 3.5 g/L of Phytagel] in a 1.5-inch diameter circle in the center of the plate to facilitate bombardment and incubated at 25°C for 4 hours in the dark. 35S-GFP DNA plasmid (35sC4PPDKsGFPTYG) was coated onto 0.6 μm gold nanoparticles (Bio-Rad): 1 mg of gold particles were mixed with 30 μl of DNA construct (0.17 μg/μl), 25 μl of 5.0 M CaCl_2_ and 20 μl of 0.1 M spermidine and incubated on ice for 10 min. DNA-coated gold particles were collected at 14,000 rpm for 1 min, and the pellet was rinsed with 1 mL of absolute alcohol, resuspended in 85 μl ethanol, and then immediately loaded onto the center of a macrocarrier (5 μl each) and allowed to air dry. Biolistic bombardment was performed using a PDS1000/He particle bombardment system (Bio-Rad) with a target distance of 6.0 cm and a rupture pressure of 900 PSI. After bombardment, leaves were transferred to MS solid medium and imaged at 3 and 10 days-post-bombardment.

### Digital droplet PCR (ddPCR) experiments

Genomic DNA (gDNA) was extracted from leaves via CTAB extraction modified from a previous method^8^. Briefly, 200 mg leaf tissue was ground in liquid nitrogen with a mortar and pestle, and the leaf powder was transferred into 600 μl CTAB buffer (10 g CTAB, 50 mL 1 M Tris-HCl pH 8, 20 mL 0.5M EDTA pH 8, 140 mL 5 M NaCl and 5 g PVP). The mixture was vortexed well and incubated at 65°C for 45 minutes. 600 μl chloroform: isopropanol (39:1) was added to mixture and vortexed well. The mixture was centrifuged at 18,000g for 10 minutes and the upper phase was transferred into a new microcentrifuge tube. 600 μl isopropanol was added to the new tube, incubated 5 minutes at room temperature, and then mixed softly. The mixture was centrifuged at 18,000g for 10 minutes. The supernatant (isopropanol) was removed and 100 μl 70% ethanol was added. The mixture was centrifuged at 18,000g for 10 minutes. The supernatant (ethanol) was removed as much as possible and the tube was left to dry at 37°C for 30 minutes. The gDNA pellet was resuspended in 200 μl autoclaved MilliQ water and the concentration and purity was measured by Nanodrop. All gDNA samples were digested overnight with HindIII-HF in CutSmart buffer. 2 μg gDNA was digested with 20U enzyme in a 50 μl reaction volume for 16 hours at 37°C. Note that the restriction enzyme was selected so as not cut inside the reference or target gene.

ddPCR was performed via probe chemistry in a duplex assay for reference EF1 and target GFP genes. The GFP probe (5’-TGCCGTCCTCCTTGAAGTCG-3’) was labeled with HEX at 5’, Iowa Black Hole Quencher at the 3’ end and with an internal ZEN quencher 9 nucleotides away from the 5’ end. The EF1 probe (5’-AGGTCTACCAACCTTGACTGGT-3’) was labeled with FAM at the 5’end, with Iowa Black Hole Quencher at the 3’ end, and with internal ZEN quencher 9 nucleotides away from the 5’ end. Primers used for GFP gene: 5’-GACTTCTTCAAGTCCGCCAT-3’ and 5’-CGTTGTGGCTGTTGTAGTTG-3’, primers used for EF1 gene: 5’-TCCAAGGCTAGGTATGATGA-3’ and 5’-GGGCTCATTAATCTGGTCAA-3’. 20X probe-primer mixes (18 μM PCR primers (each), 5 μM probe) were prepared for both genes.

ddPCR reaction mixes were prepared according to the instructions in ddPCR Supermix for Probes (No dUTP) #1863024 kit. For each sample, we prepared 10 wells, each containing 100 ng digested gDNA, so that a total of 1μg DNA was screened for transgene integration for each sample. Droplets were generated with a QX200 droplet generator right after the ddPCR reaction mixes were prepared. 20 μL of each sample mastermix was transferred to the sample row and 70 μL droplet generation oil was transferred to the oil row in the droplet generation cartridge. After the droplets were generated, 40 μL of droplets were transferred to a new 96-well plate and the plate was sealed for 5 s at 180 °C in plate sealer. The PCR was run in a deep-well thermal cycler with the following PCR program: enzyme activation 95°C 10 min, denaturation 94°C 30 sec (40X), annealing/extension 60°C 1 min (40X), stabilization 98°C 10 min, and hold 4°C. The fluorescence of the droplets was measured 4 hours after PCR (kept in the dark and at 4°C) with a QX200 droplet reader, and the results were analyzed with the Bio-Rad Quantasoft Pro Software.

### Plant toxicity analysis

To test for plant stress and toxicity, the expression level of an oxidative stress gene (NbRbohB)^9^ in *Nb* leaves was measured through qPCR with the following primers: forward 5’-TTTCTCTGAGGTTTGCCAGCCACCACCTAA-3’ and reverse 5’-GCCTTCATGTTGTTGACAATGTCTTTAACA-3’. EF1 was again measured as a housekeeping gene with the same primer set as described above. An annealing temperature of 60°C was used for qPCR, which we ran for 40 cycles, and the ddCt method was used to obtain the normalized NbRbohB expression-fold change with respect to the EF1 housekeeping gene and control sample.

As an additional toxicity assay, Fv/Fm ratios^10^ of infiltrated *Nb* leaves were measured with an Imaging-PAM Maxi fluorimeter (Walz). A singular leaf was infiltrated from the abaxial surface, in three distinct locations within the same leaf, with buffer (0.1 M NaCl), 1 mg/L DNA-CNTs, or 10% SDS (positive control for toxicity). The fourth quadrant of the leaf was left unperturbed. The triply-infiltrated leaf was subsequently incubated for 24 hours without further perturbation. Subsequently, the infiltrated leaf was dark-adapted for 15-30 minutes and chlorophyll fluorescence-related parameters were measured with the Imaging-PAM Maxi fluorimeter to calculate the Fv/Fm ratio, commonly used to test for plant stress.

### Protoplast isolation from *Eruca sativa* leaves

Protoplasts were isolated from arugula leaves as described by Yoo *et al*.^2^ with some modifications. Briefly, thinly cut arugula leaf strips were immersed in 20 mL of enzyme solution (consisting of cellulase and macerozyme), vacuum infiltrated for an hour in the dark using a desiccator, and further incubated at 37°C for 3 hours in the dark without stirring. Undigested leaf tissue was removed by filtration with a 75 μm nylon mesh, and the flow-through was centrifuged at 200 g for 3 min to pellet the protoplasts in a round bottom tube. Pelleted protoplasts were resuspended in 0.4 M mannitol solution (containing 15 mM MgCl_2_ and 4 mM MES) with a pH of 5.7, which has similar osmolarity and pH to the protoplasts. Isolated protoplasts can be kept viable on ice for over 24 h; however, we used only freshly isolated protoplasts for all gene expression studies.

### ssDNA-SWCNTs internalization by protoplasts

ssDNA-SWCNT conjugates were prepared by the same protocol described in the section titled **SDS-CNT, siRNA-CNT, and ssDNA-CNT preparation**. For protoplast internalization experiments, the oligonucleotide (GT)6 was used to prepare ssDNA-SWCNTs. Healthy protoplasts were isolated from 1-month old arugula plants. Once extracted, the protoplasts were stored in the dark at 4°C until internalization experiments were ready to be performed.

200 μL of the 3×10^5^ cells/mL protoplast suspension was mixed with 48 μL of 15.5 mg/L ssDNA-SWCNT. The samples were tapped lightly every 15 minutes to encourage mixing and prevent protoplasts from settling at the bottom of the tube. Samples were incubated for 9 hours at 4°C. The supernatant containing excess free ssDNA-SWCNT was removed without disturbing the protoplast pellet. The protoplasts were immediately resuspended in 200 μL of MMG solution. 200 μL of the protoplast suspension was transferred to a poly-L-lysine coated microwell dish and the protoplasts were allowed to settle at room temperature for 1 hour.

Immediately before imaging, 150 μL of the sample was removed from the microwell dish. All images were captured using a custom built near-infrared inverted microscope equipped with a Raptor Ninox VIS-SWIR 640 camera. Brightfield images were captured with a 100 ms exposure time. Near-infrared images were captured using a 720 nm excitation laser with a 200 ms exposure time and with a 1070 nm long pass filter.

### Protoplast transformation with DNA-CNTs prepared via dialysis

100 μL (~2×10^4^) of isolated protoplasts in mannitol solution were added to 10 μg DNA containing DNA-CNT dialysis solution (see section titled **Direct adsorption of DNA onto CNTs via dialysis**), or for the control sample only 10 μg plasmid DNA and mixed well by gently tapping the tube. The mixture was incubated at room temperature for 1 h, and subsequently centrifuged at 200g for 3 min to pellet protoplasts. Protoplasts were resuspended in 1 mL of 0.5 M mannitol solution (containing 4mM MES and 20 mM KCl at pH 5.7) in a non-culture treated 6 well-plate (Corning) for 24 hours in the dark. Protoplasts settled at the bottom of the well plate. Fluorescence microscopy was performed through the well-plate to image the protoplasts and to measure GFP expression for quantification of transformation efficiency.

### Quantitative Western blot experiments and data analysis

Infiltrated plant leaves were harvested after 48 h and grounded in liquid nitrogen to get dry frozen powders. The frozen powders were transferred to a tube with preprepared lysis buffer containing 10 mM Tris/HCl (pH 7.5), 150 mM NaCl, 1 mM EDTA, 0.1% NP-40, 5% glycerol, and 1% Cocktail. After lysis at 4°C overnight, the tube was centrifuged at 10,000 rpm for 15 minutes and the supernatant containing whole proteins was collected to a new tube. After quantification of the total extracted proteins by Pierce 660 nm Protein Assay (Thermo, Prod# 22660), 0.5 μg of normalized total proteins from each sample were analyzed by 12% SDS–PAGE and blotted to PVDF membrane. The membrane was blocked for 1 hour using 7.5% BSA in PBST (PBS containing 0.1 % Tween20) buffer and incubated overnight at 4°C with the primary GFP antibody as required (1:2000 dilution, Abcam, ab290). After extensive washing, the corresponding protein bands were probed with a goat anti-rabbit horseradish peroxidase-conjugated antibody (1:5000 dilution, Abcam, ab205718) for 30 min. The membrane was then developed by incubation with chemiluminescence (Amersham ECL prime kit) plus and imaged by ChemiDoc™ XRS+ System (BIORAD). The intensity of GFP bands were quantified with ImageJ software.

### Quantitative PCR (qPCR) experiments for gene silencing

Two-step qPCR was performed to quantify GFP gene silencing in transgenic *Nb* plants with the following commercially-available kits: RNeasy plant mini kit (QIAGEN) for total RNA extraction from leaves, iScript cDNA synthesis kit (Bio-Rad) to reverse transcribe total RNA into cDNA, and PowerUp SYBR green master mix (Applied Biosystems) for qPCR. The target gene in our qPCR was mGFP5 (GFP transgene inserted into *Nb*), and EF1 (elongation factor 1) as our housekeeping (reference) gene. Primers for these genes were ordered from IDT. For mGFP5, primers used are: forward 5’-AGTGGAGAGGGTGAAGGTGATG-3’ and reverse: 5’-GCATTGAACACCATAAGAGAAAGTAGTG-3’. Primers for EF1 are: forward: 5’-TGGTGTCCTCAAGCCTGGTATGGTTGT-3’ and reverse: 5’-ACGCTTGAGATCCTTAACCGCAACATTCTT-3’. Annealing temperature of 60°C was used for qPCR, which we ran for 40 cycles.

qPCR data was analyzed by the ddCt method^6^ to obtain the normalized GFP gene expression-fold change with respect to the EF1 housekeeping gene and control sample. For each sample, qPCR was performed as 3 technical replicates (3 reactions from the same isolated RNA batch), and the entire experiment consisting of independent infiltrations and RNA extractions from different plants was repeated 3 times (3 biological replicates).

### Single molecule TIRF to image RNA protection by CNTs

The antisense siRNA strand from our GFP gene silencing experiments was utilized in this assay: 5’-r(UUCCGUAUGUUGCAUCACC)d(TT)-3’. 10 μM 5’ labelled Cy3-RNA was added to an equal mass of SWCNTs. The RNA-CNT suspension and removal of unbound RNA followed the same protocol as described in **SDS-CNT, siRNA-CNT, and ssDNA-CNT preparation**. The positive control comprised of the same sequence that was 5’ Cy3 labeled, and 3’ biotin labeled. 6-channel μ-slides (ibidi, μ-Slide VI 0.5 Glass Bottom) were initially washed by pipetting 100 μL of 100 mM sterile NaCl solutions into one reservoir and removing 60 μL the other end, leaving just enough solution to fully wet the channel. Each subsequent step involved depositing the desired solution volume into the reservoir and removing the equivalent volume from the other end of the channel. Slide preparation was done as described by Kruss and Landry *et al*.^11^ with some modifications. Briefly, 50 μL of 0.25 mg/mL BSA-Biotin was added to coat the surface of the glass slide for 5 minutes. Next, 50 μL of 0.05 mg/mL NeutrAvidin was added, followed by 50 μL of 1.0 mg/L RNA-SWCNT, which non-specifically adsorbs to NeutrAvidin. For the positive control, 50 μL of 200 pM biotinylated Cy3-RNA was added in place of ssRNA-SWCNT. The addition of each component comprised of a 5-minute incubation period, followed by flushing the channel with 50 μL of NaCl solution to remove specimens that were not surface-immobilized. Each channel was exposed to 50 μL of 10 μg/mL RNase A for 15 minutes at room temperature. The reaction was stopped by rinsing the channel with 50 μL NaCl solution. Slides were imaged with Zeiss ELYRA PS.1 microscope immediately following incubation with RNase A.

### Statistics and data analysis

#### AFM height data

N = 10 replicates are measurements of heights of different SWCNTs within the same SWCNT suspension. Data are expressed as each measurement together with error bars indicating standard deviation. Significance is measured with one-way ANOVA with Tukey’s multiple comparisons test. F = 885.9, COOH-SWCNT vs. PEI-SWCNT *P* < 0.0001 and PEI-SWCNT vs. DNA-PEI-SWCNT *P* < 0.0001.

#### Zeta potential data

N = 5 replicates are zeta potential measurements of the same SWCNT suspension. Data are expressed as each measurement together with error bars indicating standard deviation. Significance is measured with one-way ANOVA with Tukey’s multiple comparisons test. F = 753.2, COOH-SWCNT vs. PEI-SWCNT *P* < 0.0001 and PEI-SWCNT vs. DNA-PEI-SWCNT *P* = 0.0191.

#### Leaf GFP expression data

N = 3 replicates are independent experiments; 3 separate leaves infiltrated per sample and imaged. Each independent sample replicate contains 15 technical replicates (15 measurement from the same leaf). Confocal images reported in Figure 2b, c and 3a, d, e are representative images chosen from the results obtained in 3 independent experiments. Data are expressed as each mean from the 3-independent experiments together with error bars indicating standard error of the mean. Significance is measured with one-way ANOVA with Tukey’s multiple comparisons test. In Figure 2e, F = 22.33, Dialysis vs. electrostatic grafting samples *P* < 0.0001, and lDNA-PEI-SW vs. pDNA-PEI-SW *P* = 0.001. In Figure 3b, significance is measured with two-way ANOVA with Sidak’s multiple comparisons test. DNA-CNT Day 3 vs. Day 10 *P* = 0.0001. For qPCR results reported in Figure 3c, N = 3 replicates are independent experiments; 3 separate leaves infiltrated per sample and measured with qPCR. Each sample in each independent experiment consisted of 3 technical replicates of the qPCR reaction. Data are expressed as each mean from the 3-independent experiments together with error bars indicating standard error of the mean. Significance is measured with two-way ANOVA with Sidak’s multiple comparisons test. DNA-CNT Day 3 vs. Day 10 *P* = 0.0003. In Figure 3f, significance is measured with two-way ANOVA with Sidak’s multiple comparisons test. *Agrobacterium* Day 3 vs. Day 10 *P* = 0.012. For qPCR results reported in Figure 3g, N = 3 replicates are independent experiments; 3 separate leaves infiltrated per sample and measured with qPCR. Each sample in each independent experiment consisted of 3 technical replicates of the qPCR reaction. Data are expressed as each mean from the 3-independent experiments together with error bars indicating standard error of the mean. Significance is measured with two-way ANOVA with Sidak’s multiple comparisons test. *Agrobacterium* Day 3 vs. Day 10 *P* = 0.0028.

#### Protoplast GFP expression data

N = 5 replicates are independent experiments; 5 separate protoplast solutions are incubated with samples and imaged with fluorescence microscopy. Images reported in Figure 4b and c are representative images chosen from the results obtained in 5 independent experiments. % transformation efficiency data are expressed as each mean from the 5-independent experiments together with error bars indicating standard error of the mean. Significance is measured with one-way ANOVA with Tukey’s multiple comparisons test. F = 123.5, Buffer vs. DNA-CNT *P* < 0.0001, and Free DNA vs. DNA-CNT *P* < 0.0001.

#### siRNA silencing data

Western blot experiment has N = 3 replicates that are independent experiments, and Figure 5b denotes the results from a representative blot. Relative GFP amount data determined from the Western blot are expressed as mean from the 3-independent experiments together with error bars indicating standard error of the mean. Significance is measured with one-way ANOVA with Tukey’s multiple comparisons test. F = 54.65, NT-CNT vs. siRNA-CNT *P* = 0.0001. For GFP mRNA fold change experiments in Figure 5c, N = 3 replicates are independent experiments, starting with RNA extraction from different leaves until the qPCR amplifications. Each qPCR reaction in 3 independent experiment is done in triplicate. GFP mRNA fold change data are expressed as each mean from the 3-independent experiments together with error bars indicating standard error of the mean. Significance is measured with two-way ANOVA with Sidak’s multiple comparisons test. Free siRNA vs. siRNA-CNT *P* = 0.0002, and siRNA-CNT Day 1 vs. Day 7 *P* = 0.0012.

#### smTIRF microscopy data

For each sample, N = 3 replicates are 3 channels on microscopy slide that were prepared independently. Each channel was used to obtain 30 fields of views (technical replicates). In Figure 5d, data are expressed as each mean from the 3-independent channels together with error bars indicating standard error of the mean. Significance is measured with two-tailed unpaired t-test. F = 303.7 and *P* < 0.0001.

### Data availability statement

Authors confirm that all relevant data are included in the paper and/or its supplementary information files.

## References

1. Daniell, H., Datta, R., Varma, S., Gray, S. & Lee, S.-B. Containment of herbicide resistance through genetic engineering of the chloroplast genome. Nat Biotech 16, 345–348 (1998).

2. Liu, Y. et al. A gene cluster encoding lectin receptor kinases confers broad-spectrum and durable insect resistance in rice. Nat Biotech 33, 301–305, doi:10.1038/nbt.3069 (2015).

3. Li, T., Liu, B., Spalding, M. H., Weeks, D. P. & Yang, B. High-efficiency TALEN-based gene editing produces disease-resistant rice. Nat Biotech 30, 390–392, doi:10.1038/nbt.2199 (2012).

4. Zhang, G. et al. Overexpression of the soybean GmERF3 gene, an AP2/ERF type transcription factor for increased tolerances to salt, drought, and diseases in transgenic tobacco. Journal of Experimental Botany 60, 3781–3796, doi:10.1093/jxb/erp214 (2009).

5. Chen, Q. & Lai, H. Gene delivery into plant cells for recombinant protein production. Biomed Res Int 2015, 932161, doi:10.1155/2015/932161 (2015).

6. Himmel, M. E. et al. Biomass Recalcitrance: Engineering Plants and Enzymes for Biofuels Production. Science 315, 804 (2007).

7. Tufekcioglu, A., Raich, J., Isenhart, T. & Schultz, R. Biomass, carbon and nitrogen dynamics of multi-species riparian buffers within an agricultural watershed in Iowa, USA. Agroforestry Systems 57, 187–198 (2003).

8. Altpeter, F. et al. Advancing Crop Transformation in the Era of Genome Editing. Plant Cell 28, 1510–1520, doi:10.1105/tpc.16.00196 (2016).

9. Herrera-Estrella, L., Depicker, A., Van Montagu, M. & Schell, J. Expression of chimaeric genes transferred into plant cells using a Ti-plasmid-derived vector. Nature 303, 209–213 (1983).

10. Baltes, N. J., Gil-Humanes, J. & Voytas, D. F. Genome Engineering and Agriculture: Opportunities and Challenges. Progress in Molecular Biology and Translational Science (2017).

11. Klein, T. M., Wolf, E., Wu, R. & Sanford, J. High-velocity microprojectiles for delivering nucleic acids into living cells. Nature 327, 70–73 (1987).

12. Song, S., Hao, Y., Yang, X., Patra, P. & Chen, J. Using gold nanoparticles as delivery vehicles for targeted delivery of chemotherapy drug fludarabine phosphate to treat hematological cancers. Journal of nanoscience and nanotechnology 16, 2582–2586 (2016).

13. Mizrachi, A. et al. Tumour-specific PI3K inhibition via nanoparticle-targeted delivery in head and neck squamous cell carcinoma. Nature Communications 8, 14292 (2017).

14. Demirer, G. S. & Landry, M. P. Delivering Genes to Plants. CHEMICAL ENGINEERING PROGRESS 113, 40–45 (2017).

15. Wong, M. H. et al. Lipid exchange envelope penetration (LEEP) of nanoparticles for plant engineering: A universal localization mechanism. Nano letters 16, 1161–1172 (2016).

16. Giraldo, J. P. et al. Plant nanobionics approach to augment photosynthesis and biochemical sensing. Nature materials 13, 400–408 (2014).

17. Wu, Y., Phillips, J. A., Liu, H., Yang, R. & Tan, W. Carbon nanotubes protect DNA strands during cellular delivery. ACS nano 2, 2023–2028 (2008).

18. Zheng, M. et al. DNA-assisted dispersion and separation of carbon nanotubes. Nature materials 2, 338 (2003).

19. Wang, H. et al. High-yield sorting of small-diameter carbon nanotubes for solar cells and transistors. ACS nano 8, 2609–2617 (2014).

20. Liu, Q. et al. Carbon nanotubes as molecular transporters for walled plant cells. Nano letters 9, 1007–1010 (2009).

21. Serag, M. F. et al. Trafficking and subcellular localization of multiwalled carbon nanotubes in plant cells. ACS nano 5, 493–499 (2010).

22. Wong, M. H. et al. Nitroaromatic detection and infrared communication from wild-type plants using plant nanobionics. Nature materials 16, 264–272 (2017).

23. Choi, J. H. & Strano, M. S. Solvatochromism in single-walled carbon nanotubes. Applied Physics Letters 90, 223114 (2007).

24. Alidori, S. et al. Deploying RNA and DNA with functionalized carbon nanotubes. The Journal of Physical Chemistry C 117, 5982–5992 (2013).

25. Boehr, D. D., Farley, A. R., Wright, G. D. & Cox, J. R. Analysis of the π-π stacking interactions between the aminoglycoside antibiotic kinase APH (3 ’)-IIIa and its nucleotide ligands. Chemistry & biology 9, 1209–1217 (2002).

26. Volkov, A. & Coppens, P. Calculation of electrostatic interaction energies in molecular dimers from atomic multipole moments obtained by different methods of electron density partitioning. Journal of Computational Chemistry 25, 921–934, doi:10.1002/jcc.20023 (2004).

27. Balagurumoorthy, P., Adelstein, S. J. & Kassis, A. I. Method to eliminate linear DNA from mixture containing nicked circular, supercoiled, and linear plasmid DNA. Analytical biochemistry 381, 172–174 (2008).

28. Tinland, B. The integration of T-DNA into plant genomes. Trends in Plant Science 1, 178–184, doi: https://doi.org/10.1016/1360-1385(96)10020-0 (1996).

29. McDermott, G. P. et al. Multiplexed target detection using DNA-binding dye chemistry in droplet digital PCR. Anal Chem 85, 11619–11627, doi:10.1021/ac403061n (2013).

30. Collier, R. et al. Accurate measurement of transgene copy number in crop plants using droplet digital PCR. Plant J 90, 1014–1025, doi:10.1111/tpj.13517 (2017).

31. Miyaoka, Y., Mayerl, S. J., Chan, A. H. & Conklin, B. R. Detection and Quantification of HDR and NHEJ Induced by Genome Editing at Endogenous Gene Loci Using Droplet Digital PCR. Methods Mol Biol 1768, 349–362, doi:10.1007/978-1-4939-7778-9_20 (2018).

32. Glowacka, K. et al. An evaluation of new and established methods to determine T-DNA copy number and homozygosity in transgenic plants. Plant Cell Environ 39, 908–917, doi:10.1111/pce.12693 (2016).

33. Dobnik, D., Stebih, D., Blejec, A., Morisset, D. & Zel, J. Multiplex quantification of four DNA targets in one reaction with Bio-Rad droplet digital PCR system for GMO detection. Sci Rep 6, 35451, doi:10.1038/srep35451 (2016).

34. Díaz T, P., Bernal G, A. & López C, C. Transient GUS gene expression in cassava (Manihot esculenta Crantz) using Agrobacterium tumefaciens leaf infiltration. Revista MVZ Córdoba 19, 4338–4349 (2014).

35. Yoshioka, H. et al. Nicotiana benthamiana gp91phox homologs NbrbohA and NbrbohB participate in H2O2 accumulation and resistance to Phytophthora infestans. The Plant Cell 15, 706–718 (2003).

36. Van Kooten, O. & Snel, J. F. The use of chlorophyll fluorescence nomenclature in plant stress physiology. Photosynthesis research 25, 147–150 (1990).

37. Schaumberg, K. A. et al. Quantitative characterization of genetic parts and circuits for plant synthetic biology. Nature methods 13, 94 (2016).

38. Hannon, G. J. RNA interference. Nature 418, 244–251 (2002).

39. Wittrup, A. & Lieberman, J. Knocking down disease: a progress report on siRNA therapeutics. Nature reviews. Genetics 16, 543 (2015).

40. Tang, W. et al. Post-transcriptional gene silencing induced by short interfering RNAs in cultured transgenic plant cells. Genomics, proteomics & bioinformatics 2, 97–108 (2004).

41. Sullivan, A. M. et al. Mapping and dynamics of regulatory DNA and transcription factor networks in A. thaliana. Cell reports 8, 2015–2030 (2014).

42. Lau, W. & Sattely, E. S. Six enzymes from mayapple that complete the biosynthetic pathway to the etoposide aglycone. Science 349, 1224–1228 (2015).

43. Cunningham, F. J., Goh, N. S., Demirer, G. S., Matos, J. L. & Landry, M. P. Nanoparticle-Mediated Delivery towards Advancing Plant Genetic Engineering. Trends in Biotechnology, doi: https://doi.org/10.1016/j.tibtech.2018.03.009 (2018).

## References

1. Del Bonis-O’Donnell, J. T.et al. Engineering Molecular Recognition with Bio-mimetic Polymers on Single Walled Carbon Nanotubes. JoVE (Journal of Visualized Experiments), e55030–e55030 (2017).

2. Yoo, S.-D., Cho, Y.-H. & Sheen, J. Arabidopsis mesophyll protoplasts: a versatile cell system for transient gene expression analysis. Nature protocols 2, 1565 (2007).

3. Yang, M., Gao, Y., Li, H. & Adronov, A. Functionalization of multiwalled carbon nanotubes with polyamide 6 by anionic ring-opening polymerization. Carbon 45, 2327–2333 (2007).

4. Beyene, A. G., Demirer, G. S. & Landry, M. P. Nanoparticle - Templated Molecular Recognition Platforms for Detection of Biological Analytes. Current protocols in chemical biology, 197–223 (2016).

5. Ma, L. et al. Enhanced Li–S batteries using amine-functionalized carbon nanotubes in the cathode. ACS nano 10, 1050–1059 (2015).

6. Schmittgen, T. D. & Livak, K. J. Analyzing real-time PCR data by the comparative CT method. Nature protocols 3, 1101 (2008).

7. Hajdukiewicz, P., Svab, Z. & Maliga, P. The small, versatilepPZP family ofAgrobacterium binary vectors for plant transformation. Plant Molecular Biology 25, 989–994, doi:10.1007/bf00014672 (1994).

8. Murray, M. G. & Thompson, W. F. Rapid isolation of high molecular weight plant DNA. Nucleic Acids Research 8, 4321–4326, doi:10.1093/nar/8.19.4321 (1980).

9. Yoshioka, H. et al. Nicotiana benthamiana gp91phox homologs NbrbohA and NbrbohB participate in H2O2 accumulation and resistance to Phytophthora infestans. The Plant Cell 15, 706–718 (2003).

10. Van Kooten, O. & Snel, J. F. The use of chlorophyll fluorescence nomenclature in plant stress physiology. Photosynthesis research 25, 147–150 (1990).

11. Kruss, S. et al. Neurotransmitter Detection Using Corona Phase Molecular Recognition on Fluorescent Single-Walled Carbon Nanotube Sensors. Journal of the American Chemical Society 136, 713–724, doi:10.1021/ja410433b (2014).

